# DE-SWAN analysis of biomarker trajectories over the life course is highly sensitive to age distribution and model parameterization

**DOI:** 10.64898/2026.09.29.754760

**Authors:** Anchen Che, Prisha Shah, Kamaryn T. Tanner, Alan A. Cohen

## Abstract

DE-SWAN is a commonly used method to study how large ensembles of omics biomarkers change across the life course. It has been used to demonstrate apparent repeated accelerations and decelerations of aging rates across the life course, including 2-3 peaks of aging-related changes. This result contradicts what is expected based on the demography of aging and known biological mechanisms. Here, we use simulations to study the robustness of DE-SWAN to the age distribution of the study sample and to choice of window size. We show that under many realistic scenarios, DE-SWAN is highly sensitive to factors unrelated to the underlying rates of change in aging biology. This calls into question the results of previous analyses that have attributed DE-SWAN “peaks” to acceleration and deceleration of aging. We conclude that the method is unreliable for studying rates of change across the life course under most circumstances.

## Introduction

Human aging is characterized by functional decline and an increased risk of mortality and morbidity (López-Otín et al., 2013; Kennedy et al., 2014). An accumulation of damage at the molecular level underlies this deterioration (Gladyshev et al., 2021), but we still do not fully understand the mechanisms of these processes. As omics technologies have become less expensive and more comprehensive, a number of studies have attempted to identify molecular changes associated with aging using omics data (Valdés et al., 2013; Lehallier et al., 2019; Rutledge et al., 2022; Shen et al., 2024).

Characterizing the rate of aging, especially on the level of biomarkers, is crucial because it can reveal the biological mechanisms linked to aging and its associated diseases. Monotonic changes often reflect cumulative damage, while non-monotonic changes may reflect abrupt state changes. Precise characterization of the rate of aging also facilitates characterization of each biomarker’s contribution to this rate, which allows biomarkers to be formulated into ‘ageing clocks’ and aids the selection of biomarkers as disease risk predictors or therapeutic targets (Rutledge et al. 2022).

Thus, Lehallier et al. (2019) a developed novel algorithm, differential expression-sliding window analysis (DE-SWAN), to detect non-linear changes in biomarker levels during aging. DE-SWAN gained popularity because it is a quick, straightforward method for assessing nonlinearity in aging biomarkers and provides easily interpretable timepoints for potential peaks in change. The algorithm typically operates on cross-sectional datasets where a large number of biomarkers are measured across many samples spanning a range of ages. At each age, DE-SWAN compares biomarker levels in left and right age windows centered at each age, counting the number of biomarkers that show statistically significant changes using the t-test. The use of multi-year windows ensures that the change identified persists over multiple years and is not just temporary. Peaks in these counts are interpreted as ages at which biomarkers change most rapidly. Applying DE-SWAN to plasma proteomics data, Lehallier et al. found that the number of significantly changing biomarker levels peaked at ages 34, 60 and 78 and thus concluded that aging accelerates and then decelerates at these ages. A subsequent study applying DE-SWAN to multiple omics reported similar waves, but at different ages, 44 and 60 (Shen et al., 2024).

These striking changes – acceleration and deceleration of the aging process at various moments of the life course – lack a biologically plausible mechanism. In fact, everything known about the aging process suggests that most aspects of aging should proceed monotonically (either linearly, or more likely, exponentially). There exist few known cases of non-monotonic patterns of change with age. And those cases generally involve a single peak, such as blood pressure or cholesterol, which rise and then fall at older ages (Arbeev et al., 2011). However, the rarity of such non-monotonic patterns, the individual variability in when such peaks occur, and the fact that DE-SWAN also reports peaks at younger ages mean that such patterns are unlikely to be the cause of these peaks. Furthermore, the ages of peak change did not fully agree between the Lehallier et al. and Shen et al. studies. These discrepancies led us to wonder whether the results might be a statistical artifact of the DE-SWAN method. In particular, DE-SWAN relies on p-value-based tests of statistical significance aggregated across large numbers of biomarkers. P-values are a function not only of effect size but also of statistical power, which might change across the age range for reasons unrelated to the actual rate of change in biomarker levels with age. Such confounding factors could also influence the selection of samples in the statistical tests, thus influencing the effect size.

Studies that have systematically examined DE-SWAN’s susceptibility and sensitivity to such influences are lacking. This simulation study tests the sensitivity of DE-SWAN to various factors, including the age distribution of the sample, the window size, and the functional forms of relationships between age and the individual biomarkers. We discovered that non-uniform age distributions, such as bimodal or skewed distributions, can produce false-positive peaks that reflect the shape of the age distribution rather than true biological changes. Additionally, the choice of window size affects the DE-SWAN results because it determines the total sample size and the skew of the age distributions within windows.

## Methods

### Simulation Overview

We performed a simulation study to systematically evaluate DE-SWAN under controlled conditions. We generated datasets with various biomarker trajectories simulating linear and non-linear biomarker patterns during aging and tested whether DE-SWAN could correctly identify such trajectories. A simulation study suited our aims better than analysis of real datasets since the true patterns and linearity of biomarker changes were known, allowing for direct assessment of the algorithm’s accuracy.

### Data Generating Mechanisms

To investigate the effect of age distribution in the dataset on the performance of the DE-SWAN algorithm, we simulated data using five different age distributions representing both extreme distributions and those typically found in cohort datasets (Figure 1, Table 1). A representative subset of the age distribution from the dataset from Lehallier et al. (2019) was used as an example of a real-world cohort.

**Table 1:** Simulated age distributions.

| Type | Description |
| --- | --- |
| Uniform | $\text{age}_i \sim U(18, 95)$ |
| Normal | $\text{age}_i \sim N\left(\mu = \frac{(95 + 18)}{2}, \sigma = \frac{(95 - 18)}{6}\right)$ |
| Bimodal | $\text{age}_i \sim 0.5 \cdot N(\mu_1 = 18 + 20, \sigma_1 = 8) + 0.5 \cdot N(\mu_2 = 95 - 20, \sigma_2 = 8)$ |
| Beta | $\text{age}_i \sim 18 + (95 - 18) \cdot \text{Beta}(\alpha = 1.6, \beta = 3.0)$ |
| from Lehallier et al. | Digitized from extended data figure of Lehallier et al. (2019). A Gaussian kernel density estimate (KDE) was applied to interpolate to 1-year bins. From the resulting distribution of 4,190 samples, 3,900 were randomly drawn. |

**Figure 1:**
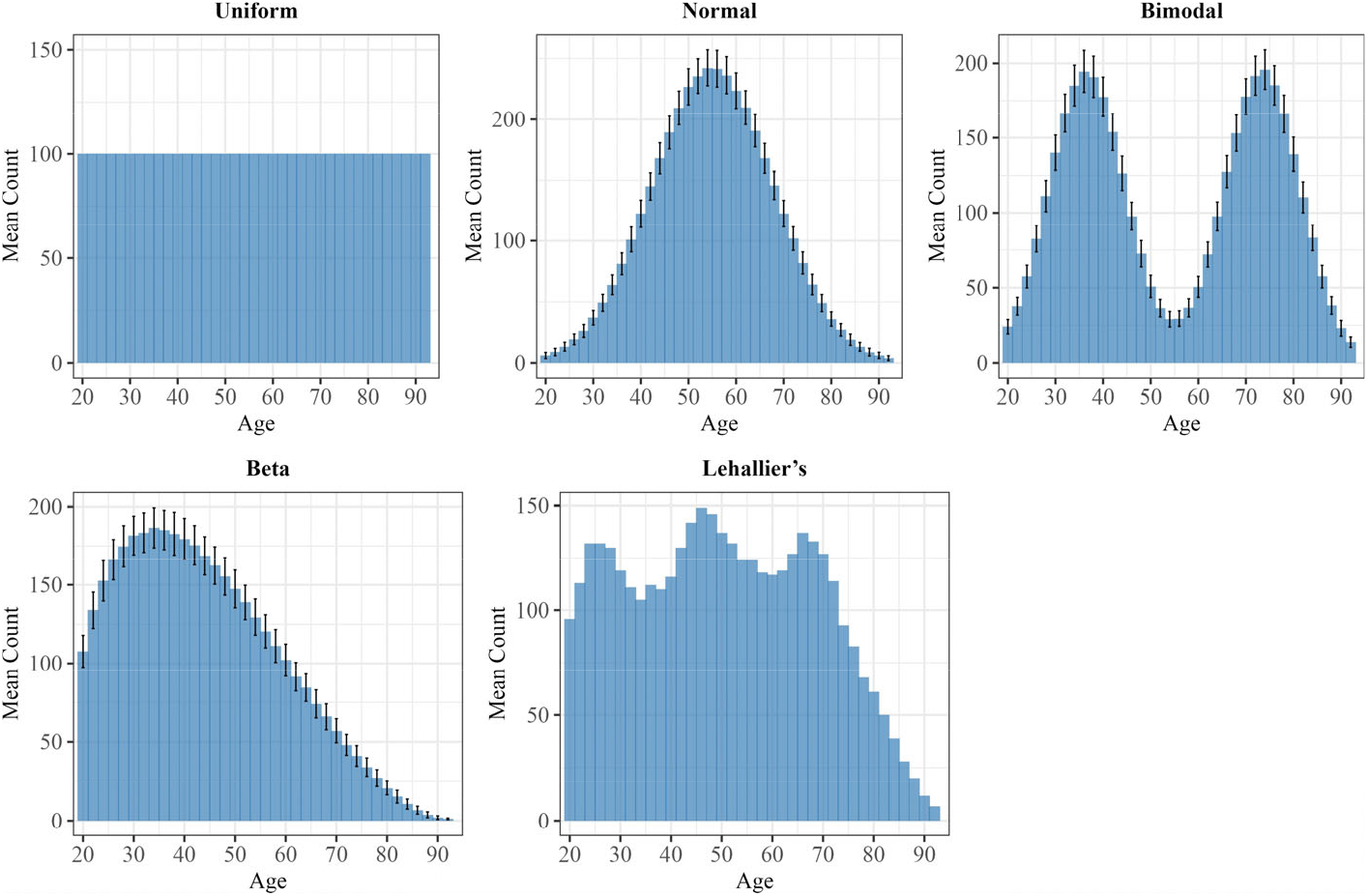
Histograms of simulated age distributions. Each panel contains the histograms for one age distribution. Normal, bimodal and beta age distributions are probabilistically generated for each simulated dataset; error bars show the standard deviation of the frequency of each age.

Uniformly distributed ages represented the reference case because the results were not affected by differing numbers of data points at different ages. The uniform age distribution was thus kept constant and deterministic. The Lehallier distribution was also kept constant, as it represented a real dataset. The normal, bimodal, and beta age distributions were regenerated for each of the 500 simulated datasets, enabling us to assess the general effect of each distribution type on DE-SWAN results. The resulting simulated ages were then used to generate biomarker trajectories. We studied the DE-SWAN algorithm using linear, exponential, piecewise-defined (linear followed by exponential), and sigmoid trajectory types (Table 2).

**Table 2:** Simulated biomarker levels.

| Trajectory shape | Description |
| --- | --- |
| Linear | $x_j = k \text{ age}_j + \varepsilon$ where $k \sim \text{LogNormal}(\mu = \log(0.03), \sigma = 0.5)$ |
| Exponential | $x_j = 0.05e^{k(\text{age}_j - 30)} + \varepsilon$ where $k \sim \text{LogNormal}(\mu = \log(0.07), \sigma = 0.1)$ |
| Linear-Exponential | For $\text{age}_j < 60$: $x_j = ke^{-20k\text{age}_j} + (1 - 60k)e^{-20k} + \varepsilon$<br>For $\text{age}_j \geq 60$: $x_j = e^{k(\text{age}_j - 80)} + \varepsilon$<br>where $k \sim \text{LogNormal}(\mu = \log(0.05), \sigma = 0.2)$ |
| Sigmoid | $x_j = m/(1 + e^{-k(\text{age}_j - 50)}) + \varepsilon$ where $m \sim \text{LogNormal}(\mu = \log(5), \sigma = 0.5)$,<br>$k \sim \text{LogNormal}(\mu = \log(0.2), \sigma = 0.5)$ |

The linear, exponential, and linear-exponential piecewise trajectories were designed to mimic the z-score trajectories of biomarker clusters shown in Figure 2d of Lehallier et al., which represent how biomarker levels change with age in the Lehallier cohort. The sigmoid trajectory, characterized by rapid changes in biomarker levels over a limited age range and relatively little change elsewhere, was included as another plausible model for the “waves of aging”, if such waves exist. The piecewise-defined trajectory was constructed by simulating the exponential trajectory and then extending it to ages below 60 such that the slope at age 60 was preserved. In other words, the trajectory for ages *<* 60 was a backward extension of the exponential curve from age 60. All trajectories included Gaussian noise *ε* ∼ N (*µ* = 0, *σ* = 1.5). All parameters of the biomarker trajectories, including the magnitude of the noise, were tuned to resemble the trajectories reported in studies such as Lehallier et al. (2019) and Shen et al. (2024).

**Figure 2:**
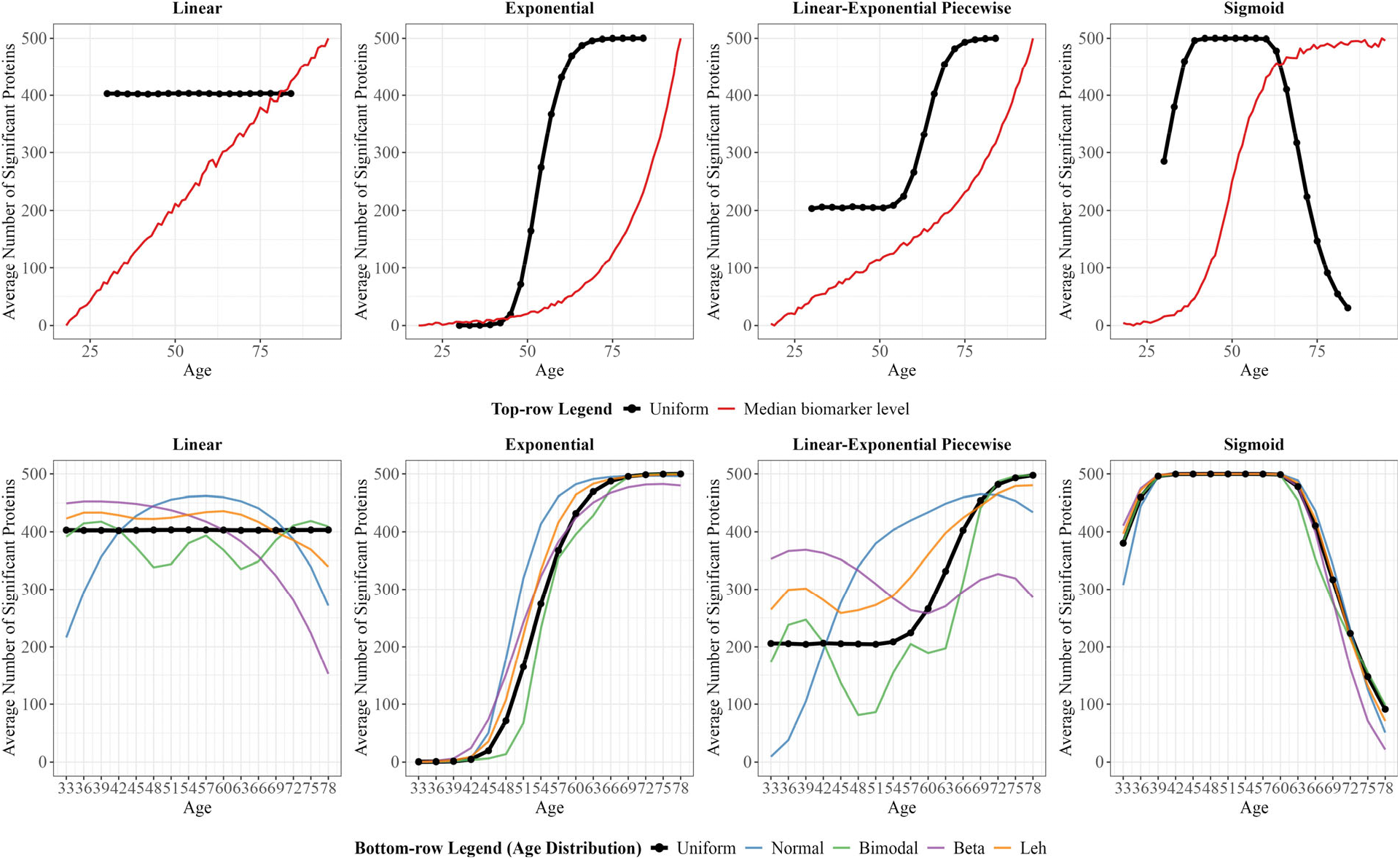
Effect of different age distributions. Each column represents a different simulated biomarker trajectory. The top row shows the DE-SWAN result (average number of significant biomarkers) for each biomarker trajectory using the uniform age distribution simulated datasets (black) and the median simulated biomarker level at each age from one simulated dataset, scaled to have a range of 0-500 (red). The bottom row shows the DE-SWAN results for each age distribution for each biomarker trajectory shape. Significant biomarkers (proteins) are those with *q <* 0.05.

We considered 20 simulation scenarios: one per combination of age distribution and biomarker trajectory. For each scenario, we simulated 500 independent datasets, each consisting of 500 biomarkers measured across 3,900 simulated individuals spanning ages 18–95 years (inclusive). The total number of individuals and the age range were chosen to approximate the distribution in Lehallier et al. (2019), which had 4,263 samples across the same age range. Each dataset represented a single-sex, single-cohort population, with no covariates used to generate biomarker levels.

### DE-SWAN

The DE-SWAN algorithm was applied to each simulated dataset. In the analysis of the effect of age distribution on performance, we applied DE-SWAN with a window size of 10 (10-year left window and 10-year right window), consistent with the implementation of Lehallier et al. (2019). In order for the windows to be fully contained in the age range of 18-95 when the window centers vary, we set the range of DE-SWAN window centers to ages 30 to 84. DE-SWAN was performed on every third year in this range as preliminary analyses indicated that performing DE-SWAN on every third year produced results consistent with those obtained when using every individual year while reducing computational burden. In a second analysis focused on the effect of window size on performance, we applied DE-SWAN with window sizes of 5, 10, and 15 years. These window sizes were chosen to balance two competing aims of the algorithm: to capture age-related changes that persist over multiple years while still attributing those changes to a relatively precise point in time. To ensure that the 15-year windows were fully contained in the age range, the window centers were restricted to every third year of age from 33 to 78. The DE-SWAN algorithm was implemented using the code provided by Lehallier et al. We calculated the average number of significant biomarkers across the 500 simulated datasets, where significance was defined as *q <* 0.05 after adjustment for multiple testing.

### Evaluation of performance

The outputs of a DE-SWAN analysis are the number of biomarkers that have significantly changed at each age (window center). In the first analysis, for a given biomarker trajectory, we compared the number of significant biomarkers found by DE-SWAN for each age distribution with the results from a uniform age distribution. Because the uniform age scenarios had an equal number of simulated individuals at each age, those results represented our reference case. For a given biomarker trajectory, differences from the uniform age scenarios highlighted the impact of age distribution on the DE-SWAN algorithm. In our second analysis, we assessed whether, for a given combination of biomarker trajectory and age distribution, DE-SWAN produced similar results across different window sizes, given that there is currently no clear guideline for selecting a window size to identify non-linear aging.

Our primary interest when comparing DE-SWAN results for an age distribution to its reference case or evaluating the similarity of results between window sizes was the increasing or decreasing trend in the number of significant biomarkers and, specifically, the ages at which the number of significant biomarkers peaked. These characteristics could have biological relevance, in particular with respect to the rate of aging. In comparison, we were less concerned about the absolute number of significant biomarkers, as this value is rarely interpreted without considering the trend, and is inevitably sensitive to window size. A larger window size is likely to yield a higher number of significantly changing biomarkers because it increases the number of observations in each window, thereby increasing statistical power. This is not necessarily problematic; DE-SWAN could still lead to similar biological conclusions if differences in window size affected the magnitude but not the trend across the life course or the location of peaks.

DE-SWAN results are typically interpreted visually, without a formal algorithm to define peak locations. Peaks are not necessarily equivalent to local maxima. In relatively smooth DE-SWAN curves, peaks can be identified as local maxima. However, in more irregular curves, a peak should be considered the maximum within a surrounding area, since numerous local maxima can otherwise obscure the true age of increased biomarker change. We also interpreted our DE-SWAN results visually. While this visual assessment introduces some subjectivity, algorithmic identification of peaks requires additional assumptions.

## Results

### Effect of age distribution on DE-SWAN results

Results of the simulations examining the effect of different age distributions on the DE-SWAN results are presented in Figure 2. For each biomarker trajectory type, we expected a different pattern in the number of significant biomarkers. These are shown in the top row of Figure 2 for data simulated with a uniform age distribution, the reference case. For example, for a sigmoid-shaped trajectory, we expected fewer biomarkers to be identified as changing at younger and older ages because the trajectories were flatter at those ages; in the middle of the age distribution where the simulated trajectories were steeper, we expected DE-SWAN to identify a greater number of changing biomarkers.

To understand how differing age distributions in the data affect the DE-SWAN algorithm, we compared results for each trajectory type for all age distributions (Figure 2-bottom row, Supplementary Figure 1). For the exponential and sigmoid biomarker trajectories, the trend and peaks in the number of changing biomarkers across ages were similar for different age distributions and similar to the reference case. For these two trajectories, at each age, the slope was either so steep that changes were detected for all age distributions, or so flat that changes were not detected in any age distribution. In real-world datasets, biomarkers with such highly pronounced trajectories may be reliably captured by DE-SWAN. In contrast, for the linear and linear-exponential biomarker trajectory shapes, the trend and peaks in the number of changing biomarkers differed markedly by age distribution. When biomarkers were simulated to change linearly, DE-SWAN correctly found an approximately constant number of changing biomarkers at all ages only for datasets with uniformly distributed ages (Figure 2). For datasets with other age distributions, the pattern in number of changing biomarkers reflected characteristics of the age distribution itself; for a normal age distribution the results peaked near the middle of the age distribution and for the bimodal age distribution peaks and valleys were seen in the results.

### Effect of window size on DE-SWAN results

Simulation results with varying DE-SWAN window sizes are presented in Figure 3. Similar to Figure 2, for the exponential and sigmoid biomarker trajectories, the trend and peaks were similar for each window size for all age distributions. However, as the window size increased, the curve shifted upwards due to the increased statistical power, as discussed in the methods section. The age at which the number of changing biomarkers reached its maximum also differed by window size. This is because in our simulation, the maximum number of changing biomarkers was 500, equal to the number of biomarkers simulated per individual. For these two trajectories, in several cases, almost all biomarkers changed significantly because the levels of all biomarkers were generated with non-constant trajectories. In a real-world dataset, more biomarkers would have constant or approximately constant trajectories, and the number of changing biomarkers would not be sustained at its maximum across ages.

**Figure 3:**
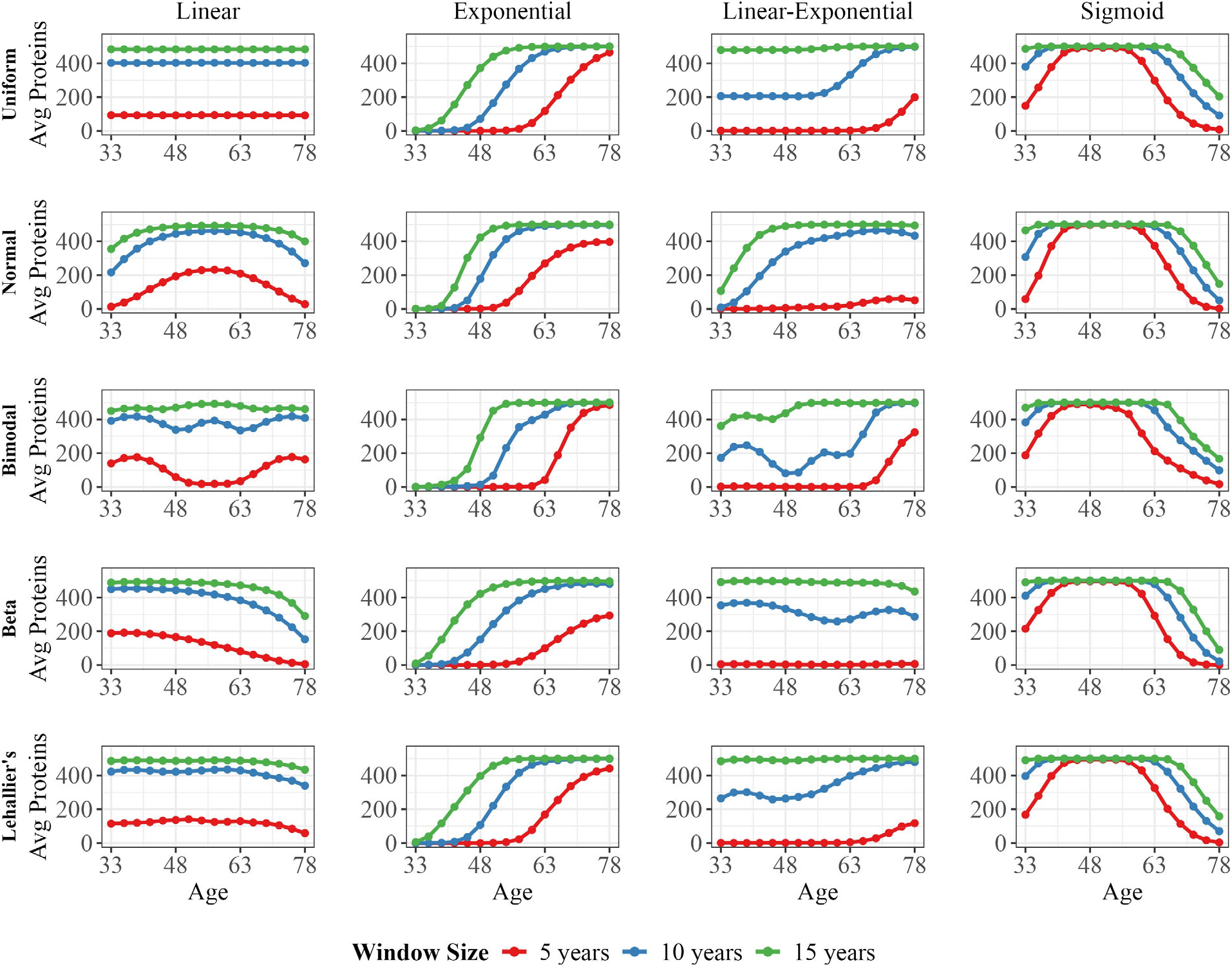
Effect of different window sizes. For each combination of age distribution (columns) and biomarker trajectory (rows), the DE-swan results (average number of significant proteins) are plotted with window sizes of 5 (red), 10 (blue) and 15 (green) years.

For the dataset with linear biomarker trajectories and a bimodal age distribution, the results from different window sizes were markedly different: a window size of 5 years produced peaks at high and low ages while a window size of 15 years produced a relatively flat pattern with a small peak in the center of the age distribution. Window size also altered the DE-SWAN results in several cases with the linear-exponential biomarker trajectory.

## Discussion

We showed that the DE-SWAN method is sensitive to both the age distribution of the sample and the choice of window size. These sensitivities do not arise from the underlying biology that DE-SWAN was meant to detect, rendering the results of DE-SWAN difficult to interpret and potentially misleading. For example, we were able to simulate multiple false peaks across age simply by changing the age distribution and/or the window size. This sensitivity can be readily explained by the factors described below. Overall, these findings suggest that DE-SWAN will rarely be an appropriate method to assess patterns in the rate of biomarker change with age.

### Explanation of age distribution effects

DE-SWAN results from datasets with the same simulated biomarker trajectories but different age distributions differed mainly for two reasons. First, a higher sampling density around an age in the age distribution of samples may cause a false-positive peak at that age. DE-SWAN uses a t-test to detect significant differences between left and right windows. Thus, a greater sample size in both windows, as occurs around a peak in the age distribution, can cause DE-SWAN to show an increased number of significantly changing biomarkers even when the difference between the mean in the left and right window is the same, as in the case of linear biomarker trajectories. In many cases, particularly for linear biomarker trajectories (excluding the bimodal age distribution), the DE-SWAN output resembles a mix of two trends: the reference result from the uniform distribution and the underlying age distribution of the sample. Peaks and troughs in the DE-SWAN curve tend to align with regions of higher or lower age density, respectively.

Second, and counterintuitively, troughs in the age distribution of samples could also lead to peaks in DE-SWAN, and vice versa. When performing DE-SWAN at an age that has fewer observations than the higher or lower ages, the age distribution within the left window would be left skewed and that within the right window would be right skewed. Since the means of the windows are further apart, the difference between their mean values increases, so t-tests are more likely to return a significant result. This effect is most noticeable in the dataset with a bimodal age distribution and a linear biomarker trajectory, where a peak at age 57 coincides with the trough of the bimodal age distribution. These two principles oppose each other, but not equally in all cases, leading to unpredictability in the DE-SWAN result in relation to age distribution.

### Explanation of window size effects

Above, we noted that the age distribution affects DE-SWAN results because (a) the sample size, and therefore the statistical power, changes with age, and (b) mean values of a biomarker change at varying distances from the window center. These two factors are also sensitive to window size: as window size increases, so does sample size and statistical power, but not equally at all window centers, depending on the functional form of the biomarkers’ relationship with age and the age distribution. Likewise, different window sizes lead to different skewness in the age distribution within both windows, influencing the difference between the averages of the two windows and, thus, the result of the t-test.

Beyond the sensitivities noted here, another challenge of DE-SWAN is the difficulty of interpreting its results. To facilitate interpretation in our simulations, each dataset contained only one type of biomarker trajectory. However, in actual datasets, the biomarkers may exhibit different trajectories, as shown in Figure 2d of Lehallier et al. (2019), and the same biomarker might also show different trajectories across individuals. DE-SWAN aggregates this heterogeneity into the number of biomarkers showing statistically significant differences at each age. Consequently, while studies of aging rates ultimately aim to draw biological conclusions from characterizations of trajectories of biomarkers, the aggregate DE-SWAN output does not reveal which trajectories underlie the observed pattern. Thus, DE-SWAN results must be presented together with characterizations of individual trajectories. However, as these trajectories provide more direct information about biomarker level changes with age, the additional information provided by DE-SWAN is unclear. Reliance on a threshold for statistical significance gives rise to another limitation on the interpretability of DE-SWAN results. Once the threshold to detect change is met, further acceleration in change will not be reflected in the DE-SWAN count.

Our findings call into question the interpretation of results by both Lehallier et al. (2091) and Shen et al (2024). For instance, the peaks reported at ages 34 and 60 in Lehallier et al. approxi m ately correspond to the two troughs in age distribution, and such troughs can lead to false peaks. Therefore, we suggest that DE-SWAN be avoided in most analyses. False peaks in DE-SWAN output could mislead researchers into believing that certain ages correspond to rapid changes in biomarker levels, potentially influencing downstream hypotheses about aging mechanisms, biomarker selection, or timing of interventions.

DE-SWAN could be applicable in cases where its results are complemented by other analyses and the dataset has specific highly controlled characteristics such as a uniform age distribution or pronounced trajectories, like those in our simulations of exponential or sigmoid trajectories. For researchers who wish to use the method, we propose several steps that may help avoid the challenges we note. First, the age distribution should be examined, and the results should be confirmed using subsamples chosen to correspond to uniform or other age distributions. Second, analyses should be repeated using varying window sizes. Results that are not robust to changing the age distribution and window size, separately or together, may be statistical artifacts. Lastly, we recommend that results be considered in light of biologically plausible mechanisms.

## Supplementary Figure

**Supplementary Figure 1:**
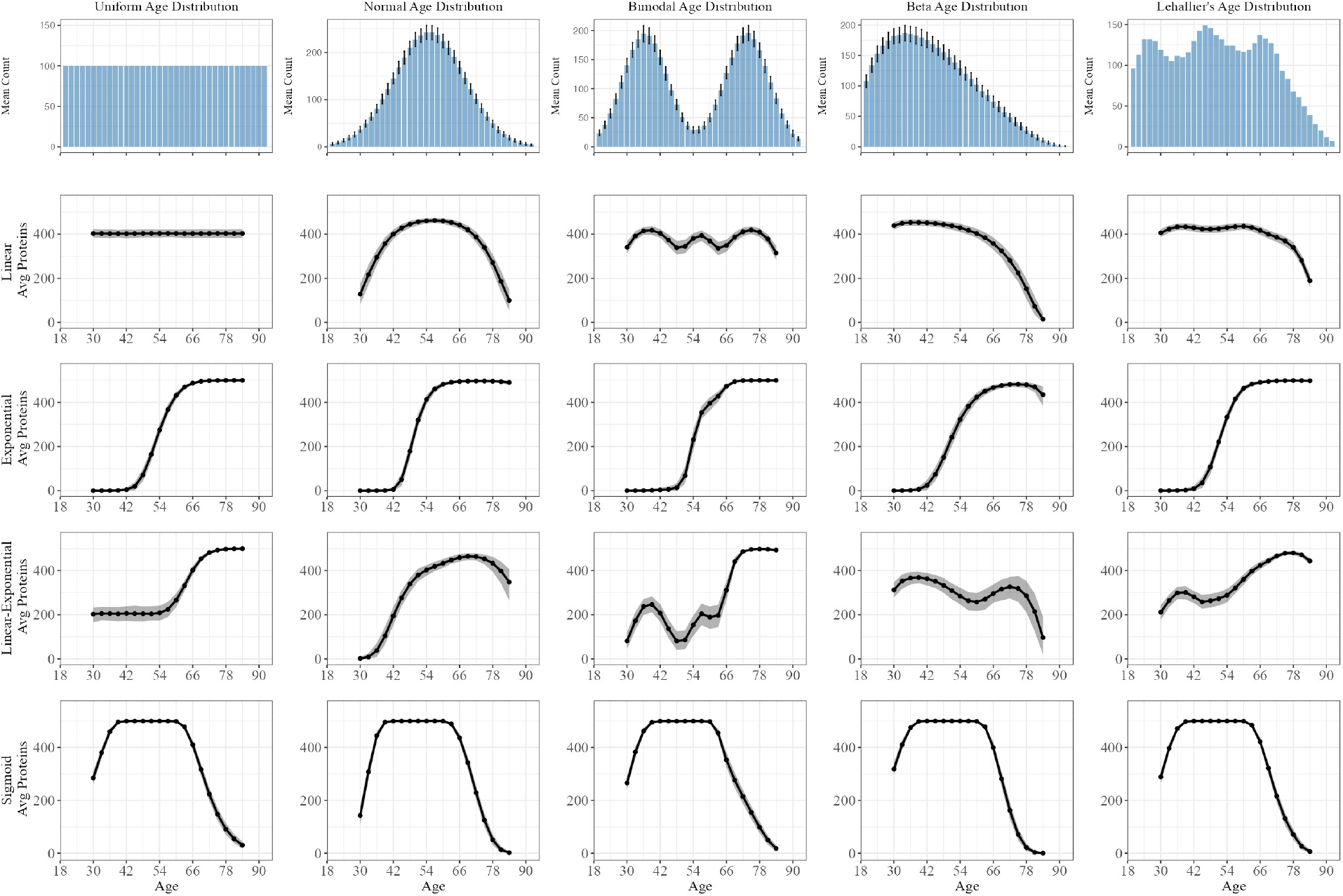
Bias due to age distribution, with confidence intervals and histogram of age distributions. Row 1 contains the histograms for the 5 age distributions. Normal, bimodal and beta age distributions are probabilistically generated for each simulated dataset and error bars show the standard deviation of the frequency of age. Rows 2-4 each represent a biomarker trajectory. For each combination of age distribution and biomarker trajectory, the median DE-SWAN result across the 500 simulated datasets is plotted with the 95% confidence interval indicated by the grey-colored shade.

## References

Arbeev, K. G., Ukraintseva, S. V., Akushevich, I., Kulminski, A. M., Arbeeva, L. S., Akushevich, L., Culminskaya, I. V., and Yashin, A. I. (2011). Age trajectories of physiological indices in relation to healthy life course. Mechanisms of Ageing and Development, 132(3):93–102.

Gladyshev, V. N., Kritchevsky, S. B., Clarke, S. G., Cuervo, A. M., Fiehn, O., de Magalhães, J. P., Mau, T., Maes, M., Moritz, R., Niedernhofer, L. J., Van Schaftingen, E., Tranah, G. J., Walsh, K., Yura, Y., Zhang, B., and Cummings, S. R. (2021). Molecular damage in aging. Nature Aging, 1(12):1096–1106.

Kennedy, B. K., Berger, S. L., Brunet, A., Campisi, J., Cuervo, A. M., Epel, E. S., Franceschi, C., Lithgow, G. J., Morimoto, R. I., Pessin, J. E., Rando, T. A., Richardson, A., Schadt, E. E., Wyss-Coray, T., and Sierra, F. (2014). Geroscience: Linking aging to chronic disease. Cell, 159(4):709–713.

Lehallier, B., Gate, D., Schaum, N., Nana′si, T., Lee, S. E., Yousef, H., Losada, P. M., Berdnik, D., Keller, A., Verghese, J., Sathyan, S., Franceschi, C., Milman, S., Barzilai, N., and Wyss-Coray, T. (2019). Undulating changes in human plasma proteome profiles across the lifespan. Nature Medicine, 25:1843–1850.

López-Otín, C., Blasco, M. A., Partridge, L., Serrano, M., and Kroemer, G. (2013). The hallmarks of aging. Cell, 153(6):1194–1217.

Rutledge, J., Oh, H., and Wyss-Coray, T. (2022). Measuring biological age using omics data. Nature Reviews Genetics, 23:715–727.

Shen, X., Wang, C., Zhou, X., Zhou, W., Hornburg, D., Wu, S., and Snyder, M. P. (2024). Nonlinear dynamics of multi-omics profiles during human aging. Nature Aging, 4:1619–1634.

Valdés, A. M., Glass, D., and Spector, T. D. (2013). Omics technologies and the study of human ageing. Nature Reviews Genetics, 14:601–607.

